# Structural Modeling of the TMPRSS Subfamily of Host Cell Proteases Reveals Potential Binding Sites

**DOI:** 10.1101/2021.06.15.448583

**Authors:** Diego E. Escalante, Austin Wang, David M. Ferguson

## Abstract

The transmembrane protease serine subfamily (TMPRSS) has at least eight members with known protein sequence: TMPRSS2, TMPRRS3, TMPRSS4, TMPRSS5, TMPRSS6, TMPRSS7, TMPRSS9, TMPRSS11, TMPRSS12 and TMPRSS13. A majority of these TMPRSS proteins have key roles in human hemostasis as well as promoting certain pathologies, including several types of cancer. In addition, TMPRSS proteins have been shown to facilitate the entrance of respiratory viruses into human cells, most notably TMPRSS2 and TMPRSS4 activate the spike protein of the SARS-CoV-2 virus. Despite the wide range of functions that these proteins have in the human body, none of them have been successfully crystallized. The lack of structural data has significantly hindered any efforts to identify potential drug candidates with high selectivity to these proteins. In this study, we present homology models for all members of the TMPRSS family including any known isoform (the homology model of TMPRSS2 is not included in this study as it has been previously published). The atomic coordinates for all homology models have been refined and equilibrated through molecular dynamic simulations. The structural data revealed potential binding sites for all TMPRSS as well as key amino acids that can be targeted for drug selectivity.

## Introduction

The Type II Transmembrane Serine Protease (TTSP) family consists of 4 different subfamilies that are differentiated by specific domains: the HAT/DESC subfamily, the Hepsin/TMPRSS subfamily, the Matriptase subfamily, and the Corin subfamily.^1^ The proteins are created as single-chain, inactive proenzymes that are activated by cleaving the basic amino acid residue, arginine or lysine, within the conserved activation motif between the pro-domain and catalytic domain.^1, 2^ Once activated, they remain associated with the membrane through disulfide bonds between the catalytic and transmembrane domains.^1^ There are currently 14 different TMPRSS proteins: TMPRSS2, TMPRSS3, TMPRSS4, TMPRSS5, TMPRSS6, TMPRSS7, TMPRSS9, TMPRSS11A, TMPRSS11B, TMPRSS11D, TMPRSS11E, TMPRSS11F, TMPRSS12, and TMPRSS13.^3^ These proteases have been found to be in multiple organs and tissues throughout the body.^4–10^ This subfamily has a diverse set of physiological functions like homeostasis and proteolytic cascades, and while not all functions have been found, their involvement in various pathogenicities is becoming apparent as shown in Table 1 below.^11^ Several inhibitors have also been identified including aprotinin, camostat, leupeptin, AEBSF, nafamostat, and gabexate.^12^

**Table 1.**
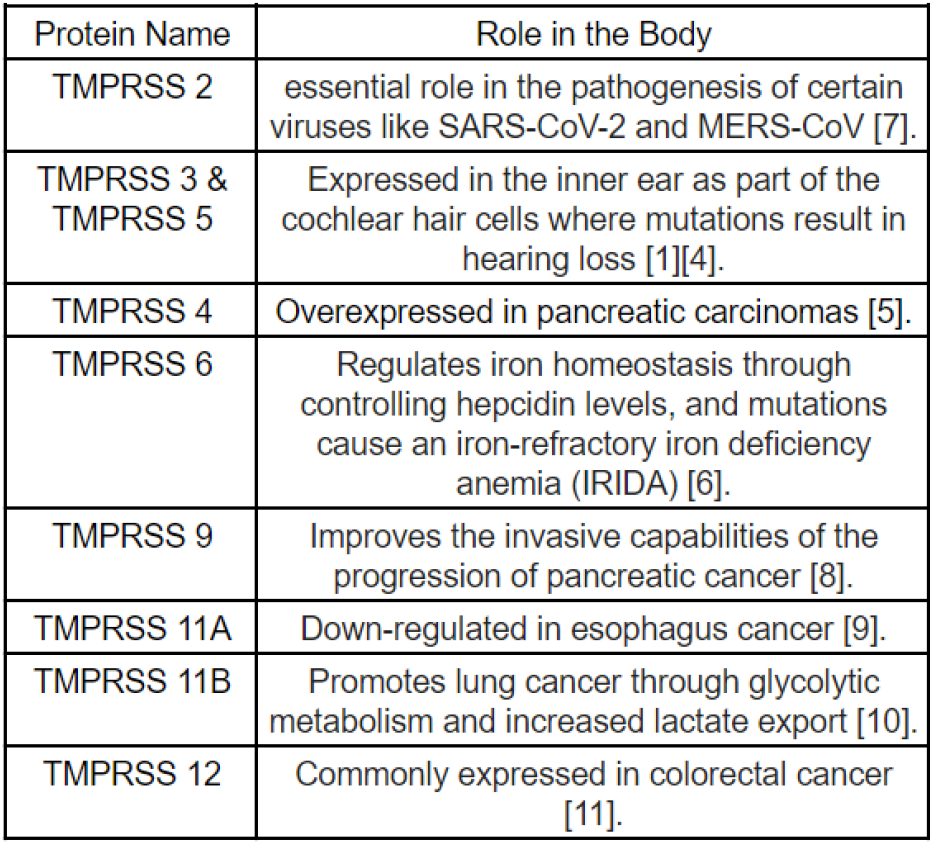
Known roles of TMPRSS proteins.

| Protein Name | Role in the Body |
| --- | --- |
| TMPRSS 2 | essential role in the pathogenesis of certain viruses like SARS-CoV-2 and MERS-CoV [7]. |
| TMPRSS 3 & TMPRSS 5 | Expressed in the inner ear as part of the cochlear hair cells where mutations result in hearing loss [1][4]. |
| TMPRSS 4 | Overexpressed in pancreatic carcinomas [5]. |
| TMPRSS 6 | Regulates iron homeostasis through controlling hepcidin levels, and mutations cause an iron-refractory iron deficiency anemia (IRIDA) [6]. |
| TMPRSS 9 | Improves the invasive capabilities of the progression of pancreatic cancer [8]. |
| TMPRSS 11A | Down-regulated in esophagus cancer [9]. |
| TMPRSS 11B | Promotes lung cancer through glycolytic metabolism and increased lactate export [10]. |
| TMPRSS 12 | Commonly expressed in colorectal cancer [11]. |

Recently, the Hepsin/TMPRSS subfamily has recently received considerable attention not only due to the potential role these enzymes play in the development of cancers but to the function TMPRSS2 plays in mediating the virulence of SARS-CoV-2. The TMPRSS2 protein and ACE2 receptor are key elements of the main pathway by which SARS-CoV-2 (and related CoV pathogens) are internalized by lung cells.^12^ The virus uses TMPRSS2 to cleave the Spike protein at a specific site (defined by Arg255 and Ile256) to activate membrane insertion.^12^ ACE2 recruits the Spike protein to the host cell surface through specific interaction with the receptor binding domain of the Spike protein.^12^ The Spike protein contains two functional subunits: the S1 subunit that allows binding of the virus to the host cell surface receptor and the S2 subunit that allows the fusion of the viral and cellular membranes.^13, 14^ In vivo studies have shown TMPRSS2-knockout mice with down-regulated TMPRSS2 show less severe lung pathology as compared to controls when exposed to SARS-CoV-2.^15, 16^ Additional work has shown that TMPRSS2 is the main processing enzyme for virus entry in lung cells.^17^

TMPRSS2 has therefore emerged as a primary target for the design and discovery of drugs for treating SARS-CoV-2 infections. An excellent starting point in the search for potent drugs is camostat. This compound is clinically approved for use in treating pancreatitis in Japan and is a known inhibitor of TMPRSS2. While camostat has shown efficacy in preventing mice infected with SARS-CoV from dying,^12^ it is a pan-trypsin-like serine protease inhibitor and is not highly selective for TMPRSS2. Similarly, there are no known or reported small molecules that are highly selective for any of the other member of the TMPRSS family. The main reason for this is the lack of crystal structures of the TMPRSS protein subfamily. Any structural information on these proteases could lead to the development of drugs more selective than the current alternatives. This study extends prior work on camostat bound to TMPRSS2 to identify common elements of recognition across the TMPRSS sub-family.

## Computational Methods

### Homology Modeling

The 14 proteins (TMPRSS2, TMPRSS3, TMPRSS4, TMPRSS5, TMPRSS6, TMPRSS7, TMPRSS9, TMPRSS11A, TMPRSS11B, TMPRSS11D, TMPRSS11E, TMPRSS11F, TMPRSS12, TMPRSS13) amino acid sequence was obtained from the UnitProt database (Gene ID: O15393, P57727, Q9NRS4, Q9H3S3, Q9DBI0, Q7RTY8, Q7Z410, Q6ZMR5, Q86T26, O60235, Q9UL52, Q6ZWK6, Q86WS5, Q9BYE2, respectively), and the crystal structures with high sequence identity, available in the Protein Databank (PDB), were retrieved through the BLASTp algorithm. The differences between proteins will be discussed in the results section. Across the 14 receptors, 25 crystal structures were used as templates to build the TMPRSS protein models. Of the 25 crystal structures, the 3 notable crystals structures were a urokinase-type plasminogen activator with a position 190 alanine mutation (1O5E), DESC1, part of the TTSP family, (2OQ5), and a human plasma kallikrein (6O1G) each with a known ligand included with each crystal structure. The 3 ligands included were 6-Chloro-2-(2-Hydroxy-Biphenyl-3-yl)-1H-indole-5-carboxamidine, benzamide, and 7SD from 1O5E, 2OQ5, and 6O1G, respectively, and were kept in the homology models created. The Pfam database was used to obtain the amino acid region corresponding to the trypsin domain for each respective TMPRSS protein. Found through the Pfam database, all the crystal structures were truncated to only their trypsin domains. The Schrodinger prime homology model suite program was used to align each sequence and identify any conserved secondary structure assignments. To construct a homology model for each TMPRSS protein, each protein had at least 5 crystal structures chosen which are listed in Supplementary Table 1. Once completed, a final consensus model was constructed by using each homology model that was built specifically for the respective protein. The final model for each TMPRSS protein is labeled as TMPRSSxhm with x being the specific name of the protein. The models were refined through a process of short molecular dynamics (MD) simulations that will be described below.

### Molecular Dynamics

All of the molecular dynamic simulation stages were completed by using the SANDER.MPI function of the AMBER 18 software package unless otherwise stated^18^. The ligand structures were pre-processed with the Antechamber package to assign AM1-BCC partial charges. The ligand and enzyme structures were processed using LEaP to assign ff15ipq^19^ or gaff^20^ and ff14SB^21^ force field parameters for the enzyme and ligand, respectively. All the complexes were submerged within a periodic box of TIP3P water with a 10Å buffer region. Each molecule was initially minimized by using the steepest descent method for 100 steps followed by 9900 steps of conjugate gradient minimization. Then, a stepwise heating procedure occurred where the system temperature was slowly ramped from 0K to 300K over 15,000 steps and then relaxed over 5,000 steps to where the average temperature was kept constant at 300k using a weak-coupling algorithm. For both heating stages, the position of all the non-solvent atoms were restrained with a harmonic potential with a force constant of 25 kcal/mol-Å. After that, a two-step procedure was performed in which the average pressure was maintained at 1bar for 20,000 steps through pressure relaxation time of 0.2ps and the Berendsen barostat. A harmonic potential with a force constant of 5 kcal/mol-Å restrained the position of all non-solvent atoms. A relaxation stage of 20,000 steps followed in which the pressure relaxation time was increased to 2ps, and the only position of the receptor C**α** was restrained by a harmonic potential with a force constant of 0.5 kcal/mol-Å. Lastly, all the production runs were carried out without any restraints and were kept at a consistent 300K temperature and 1bar pressure.

## Results

All fifteen sequences have a high enough degree of identity to template models (>40%), as shown in Figure 1. This allowed us to build homology models for all members of the TMPRSS family. All of the homology models were subject to loop refinement using the Schrodinger refinement package. The refined structures were equilibrated in a water box at 300K and 1atm as described in the methods section. All the TMPRSS structures rapidly equilibrated and their root-mean-squared-deviation plateaued at an average of 1.5-1.6Å. The equilibrated atomic coordinates are provided in the supplementary information section. A structural alignment of all equilibrated TMPRSS’ is shown in Figure 2. Most of the tertiary structure is conserved by all members of this family (i.e. the alpha-helices and beta-sheets). However, the loop found between residues 50-70 is very dissimilar, which is a result of the low sequence identity in this region. Furthermore, all TMPRSS proteins share a conserved histidine, aspartic acid, and serine that form the catalytic triad as shown in Supplementary Table 2. While the catalytic triad appears to be in different points across the amino acid sequences, the histidine, aspartic acid, and serine are all located at similar points in structural space within the trypsin region. From the structural alignment it is observed that the catalytic triad is also conserved in geometry, as shown in Figure 3.

**Figure 1.**
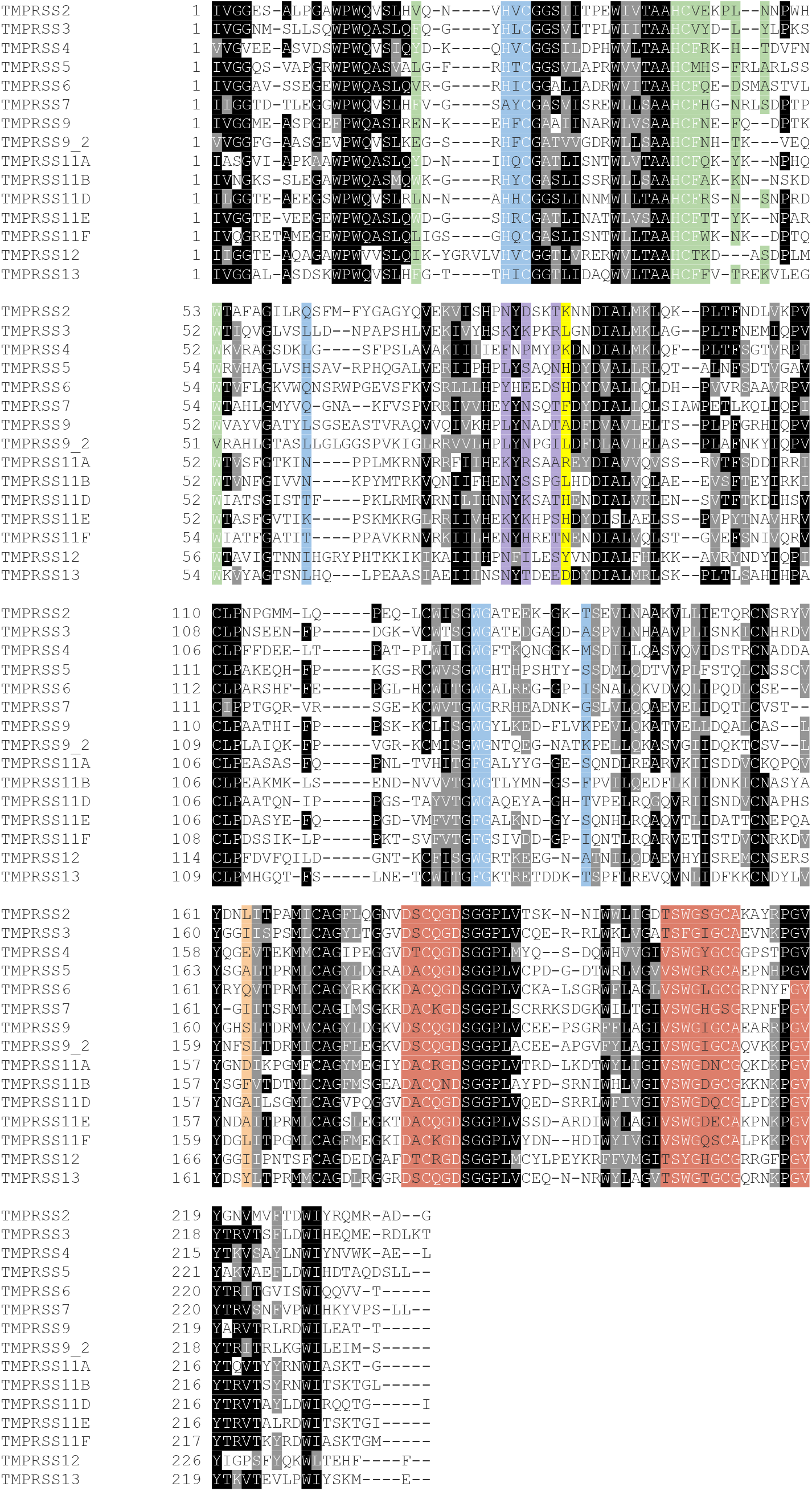
Sequence alignment of all TMPRSS family members. Residues conserved in >50% of all TMPRSS sequences are highlighted in black. From previous analysis of TMPRSS2, there are 5 potential pockets (A-E) that play a role in guiding the ligands to bind properly near the active site of TMPRSS2.^22^ Each of these components have been highlighted in Figure 1: position 87 in yellow, pocket A in red, pocket B in blue, pocket C in green, pocket D in purple, and pocket E in orange.

**Figure 2.**
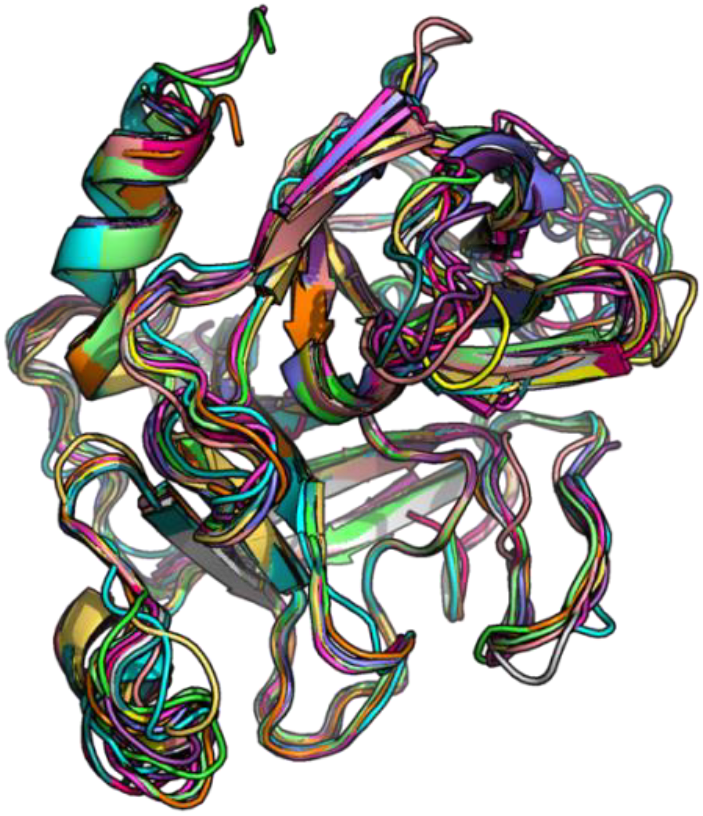
Structural alignment of refined and equilibrated TMPRSS homology models.

**Figure 3.**
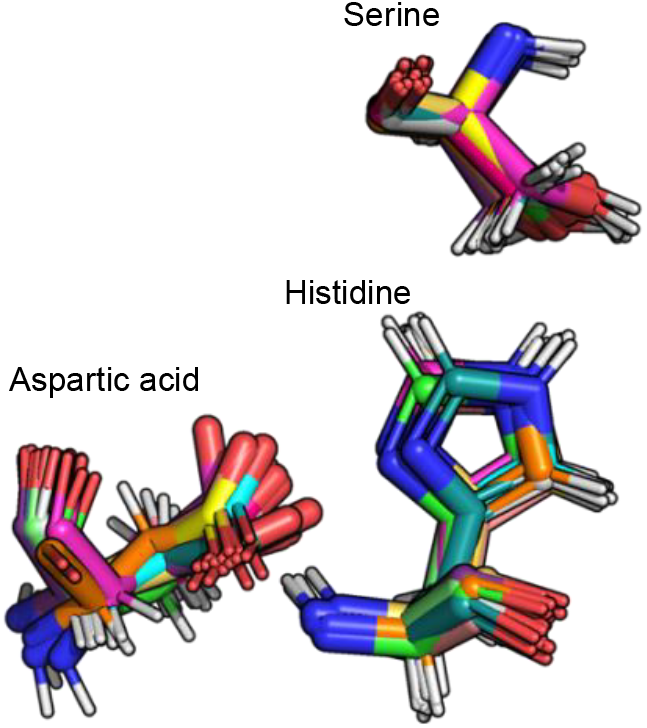
Structural alignment of the catalytic triad in all TMPRSS homology models.

### Drug Binding Site

Literature reports have shown that camostat is hydrolyzed by carboxylesterases to form the active metabolite 4-(4-guanidinobenzoyloxy) phenylacetate (GBPA), as shown in figure 4. The active metabolite GBPA has a negatively charged carboxylate group which is stabilized through the formation of a salt bridge with the lysine residue in position 87 of TMPRSS2 and simultaneously anchored via the guanidino group as shown in Figure 5.^22^ The salt bridge anchors the drug to position the scissile bond of camostat at the optimal distance for attack by the catalytic serine. The cleavage of the scissile bond by the catalytic serine results in the acylation of the enzyme, thus inhibiting TMPRSS2.

**Figure 4.**
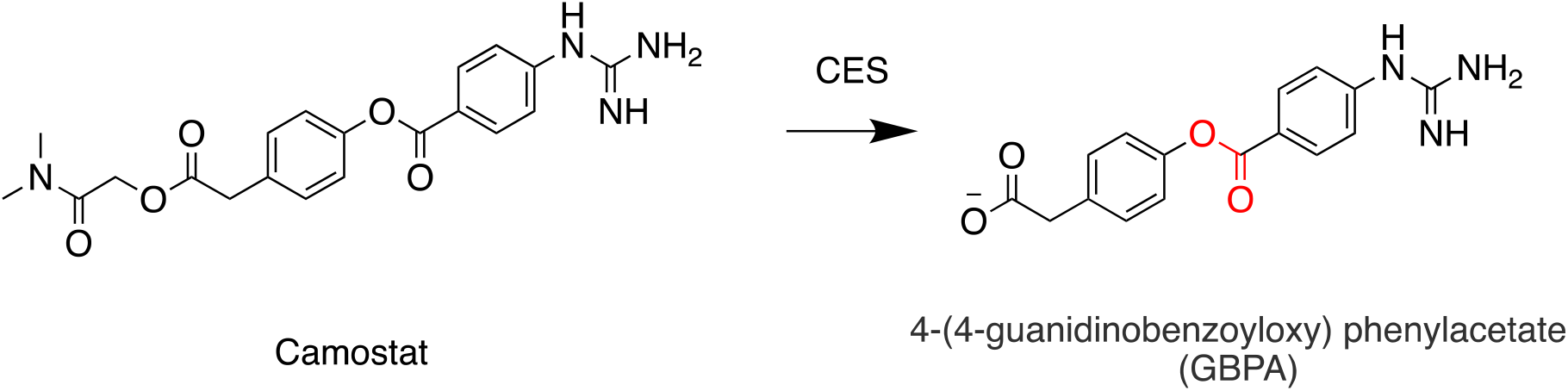
Camostat is hydrolized to the active metabolite 4-(4-guanidinobenzoyloxy) phenylacetate (GBPA) by carboxylesterases (CES). The scissile ester bond attacked by the catalytic serine of TMPRSS2 is highlighted in red.

**Figure 5.**
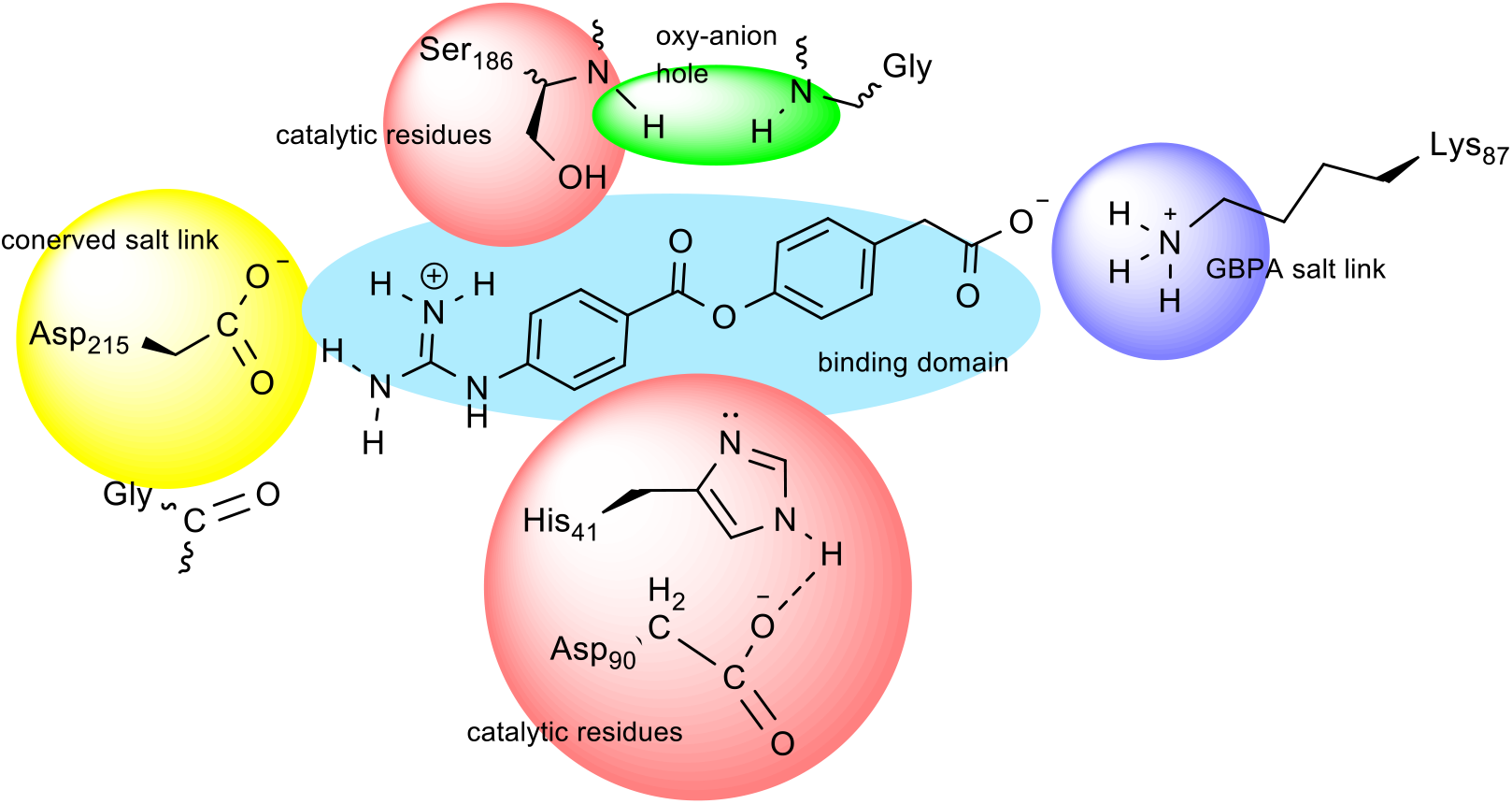
Contact interaction diagram between GBPA and TMPRSS2 active site. The anchoring residues near the active site are highlighted in yellow, the catalytic triad is highlighted in red and the lysine distal anchor is highlighted in blue.

Since K87 (K342 in Escalante et. al.^22^) has been shown to play a role in stabilizing the active metabolite of camostat,^22^ examining the other TMPRSS proteins for the amino acid in this position is crucial as it can elucidate similar stabilization mechanisms. From Figure 1, TMPRSS4 is the only other protein that conserves lysine. TMPRSS5, TMPRSS6, and TMPRSS11A maintain a positively charged amino acid through histidine and arginine, respectively. On the other hand, TMPRSS13 carries a negative charge through aspartic acid. TMPRSS11F contains the uncharged polar amino acid asparagine. TMPRSS7 and TMPRSS12 have aromatic hydrophobic side chains phenylalanine and tyrosine, respectively. All the other TMPRSS proteins contain chain hydrophobic amino acids.

### Analysis of Pockets A-E

The location of the five binding pockets identified in the TMPRSS family is shown in Figure 6. Pocket A has two critical positions that vary between TMPRSS proteins. An analysis of Figure 1 shows TMPRSS2 has an uncharged, polar amino acid residue in position 181, which is maintained in TMPRSS3, TMPRSS4, TMPRSS9, TMPRSS12, and TMPRSS13. The specific two amino acids used are serine or threonine. Other TMPRSS proteins have an alanine in this position. The second important location in pocket A is position 204 that is an uncharged, polar amino acid residue (threonine) in TMPRSS2, TMPRSS3, TMPRSS12, and TMPRSS13 while all other TMPRSS proteins have a hydrophobic side chain amino acid (e.g. valine) in that position. Pocket B shows some possible discriminant residues for ligand recognition including differences at position 25. This position is valine or isoleucine in TMPRSS2, TMPRSS3, TMPRSS4, TMPRSS6, TMPRSS12, and TMPRSS13. TMPRSS7, TMPRSS9, and TMPRSS11B contain an aromatic hydrophobic side chain. Uncharged, polar amino acids threonine and glutamine are in this position for TMPRSS5, TMPRSS11A, and TMPRSS11F whereas TMPRSS11D and TMPRSS11E have a histidine and arginine in this position, respectively. Pocket C shows significant variance across the family. At position 47, TMPRSS2 and TMPRSS3 contain a hydrophobic amino acid (i.e. leucine). For this pocket, the two isoforms of TMPRSS9 contain different amino acids at this position. TMPRSS5, one isoform of TMPRSS9, TMPRSS11A, and TMPRSS11E have aromatic hydrophobic amino acids phenylalanine and tyrosine. TMPRSS4 and TMPRSS11B have positively charged amino acids, histidine and lysine, respectively. TMPRSS6 contains the negatively charged amino acid, aspartate. TMPRSS7, the second isoform of TMPRSS9, TMPRSS11D, TMPRSS11F, and TMPRSS13 have uncharged, polar amino acids of asparagine and threonine. Lastly, TMPRSS12 has no amino acid in this matching location. Pocket D is of significant interest due to the potential interaction of GBPA and lys87 (as highlighted above). The amide of the prodrug and carboxylate of the active metabolite bind in this domain. The lysine, however, is conserved in only two proteins in this family, TMPRSS2 and TMPRSS4. While pocket E shows differences between the amino acids, interactions in this domain do not appear to play a role in ligand binding due to their locations relative to the binding pocket.

**Figure 6.**
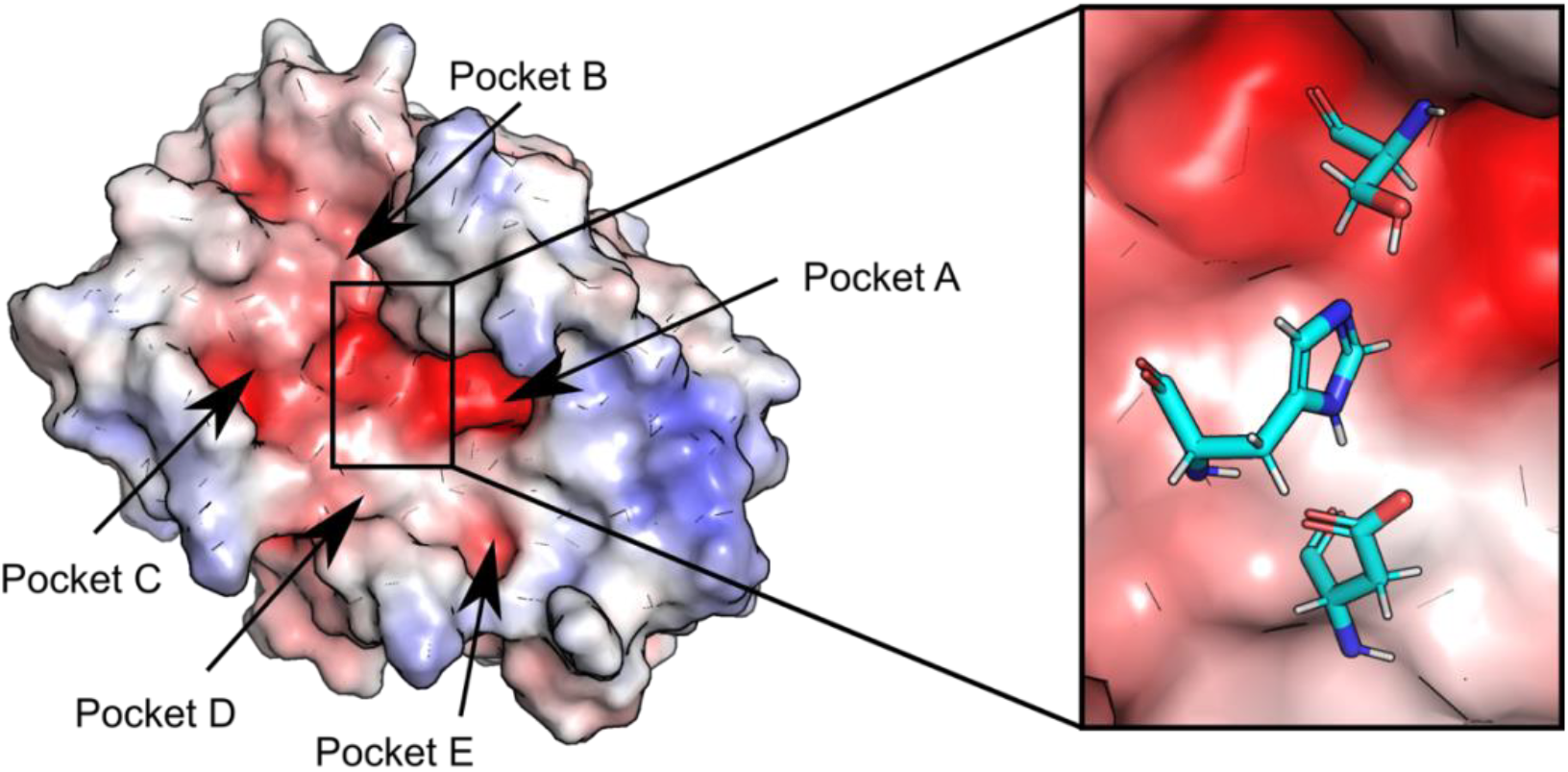
*(left)* Five binding pockets (A-E) in TMPRSS2 have been identified to be located near the active site. Pocket A corresponds to the anchor residue highlighted in yellow in **Figure 5**. The location of the active site catalytic triad is highlighted by the black rectangle. *(right)* A zoomed in view of the catalytic triad residues: serine, histidine, arginine (from top to bottom).

## Discussion

The alignments and structural analyses have shown there is significant variation across the TMPRSS family of protein to exploit in the design of selective inhibitors. Of the potential pockets and positions that influence binding, pockets A, B, C, and D have differences in the primary sequence that may drive ligand binding and selectivity. From these specific regions, differences in position 87 in pocket D may be important for ligand binding. Prior work has shown that lys87 may be a key anchor point for the carboxylate group of GBPA (the active metabolite of camostat). Only TMPRSS2 and TMPRSS4 share the conserved lysine at position 87. This is interesting because the Spike protein is known to be cleaved by both TMPRSS2 and TMPRSS4 to gain cell entry. Targeting this position may therefore provide an approach to selectively inhibit these two enzyme and reduce off target binding of camostat. Pocket A also contains an area of dense negative charges through its primary sequence. This allows for a strong ionic interaction between ligands containing positively charged amino acids and TMPRSS proteins. Because of this interaction, current ligands with large bulky positively charged regions, like nafamostat, can form strong ionic interactions with pocket A.

## Conclusion

Within the TTSP family, members of the TMPRSS subfamily have been implicated in disease progression involving cancer and infectious diseases such as COVID-19. The connection between Spike protein processing, disease progression, and TMPRSS2 activity has sparked great interest in the repurposing of drugs such as camostat and nafamostat for COVID-19 drug therapy. The alignments presented and analysis performed has shown there are significant differences in amino acid residues within the ligand binding pockets of the TMPRSS family of enzymes. Since drug selectivity and off target binding can have dramatic consequences on adverse side effects of drugs the data given here is designed to help identify unique epitopes in TMPRSS2 that can be exploited in the design of more selective agents. One of the more interesting results of this study is the identification of lys87 as a potential selectivity element for inhibiting TMPRSS2 and TMPRSS4. This lysine is only conserved in these two family members which are both enzymes that have been shown to process the SARS-CoV Spike protein. The ability to selectively block TMPRSS2,4 and avoid off target binding to other TMPRSS proteins would provide significant advantages in the development of better therapeutic agents that block protease-mediate cleavage of the Spike protein. One problem that is not easily addressed is the conserved nature of the polar-anionc region in pocket A. Drugs like nafamostat that contain a bulky cationic group are recognized across all members that display this motif. This, may in part explain the promiscuous binding of compounds showing pan-trypsin protease activity. These results suggest it may not be possible to improve the selectivity of these agents for repurposing in the design of COVID-19 agents.

## Supporting information

Homology Model Coordinates

## Supplementary Appendix

**Supplemental Figure 1.**
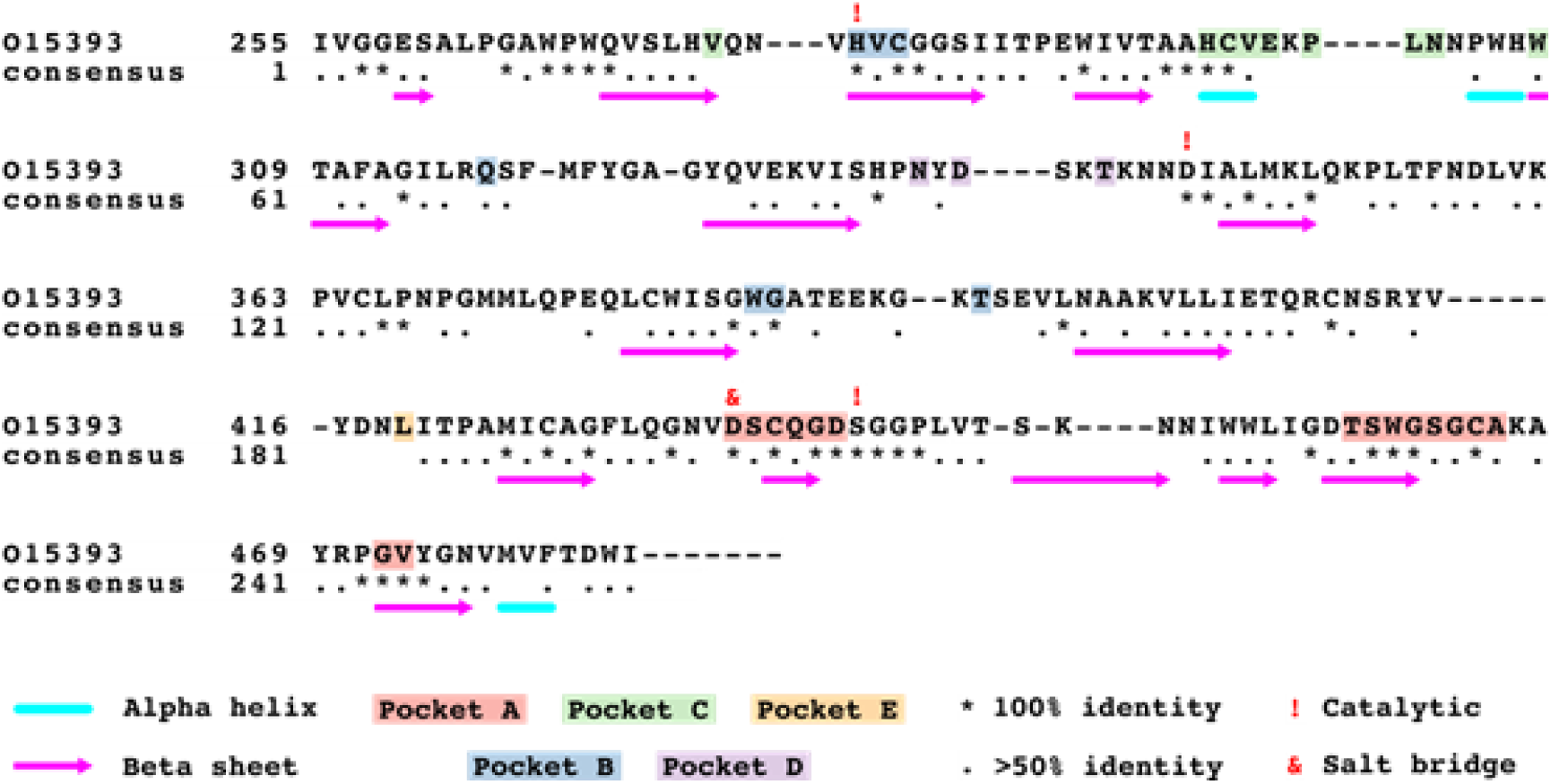
TMPRSS2 Potential Pockets for ligand binding.

**Supplementary Table 1.**
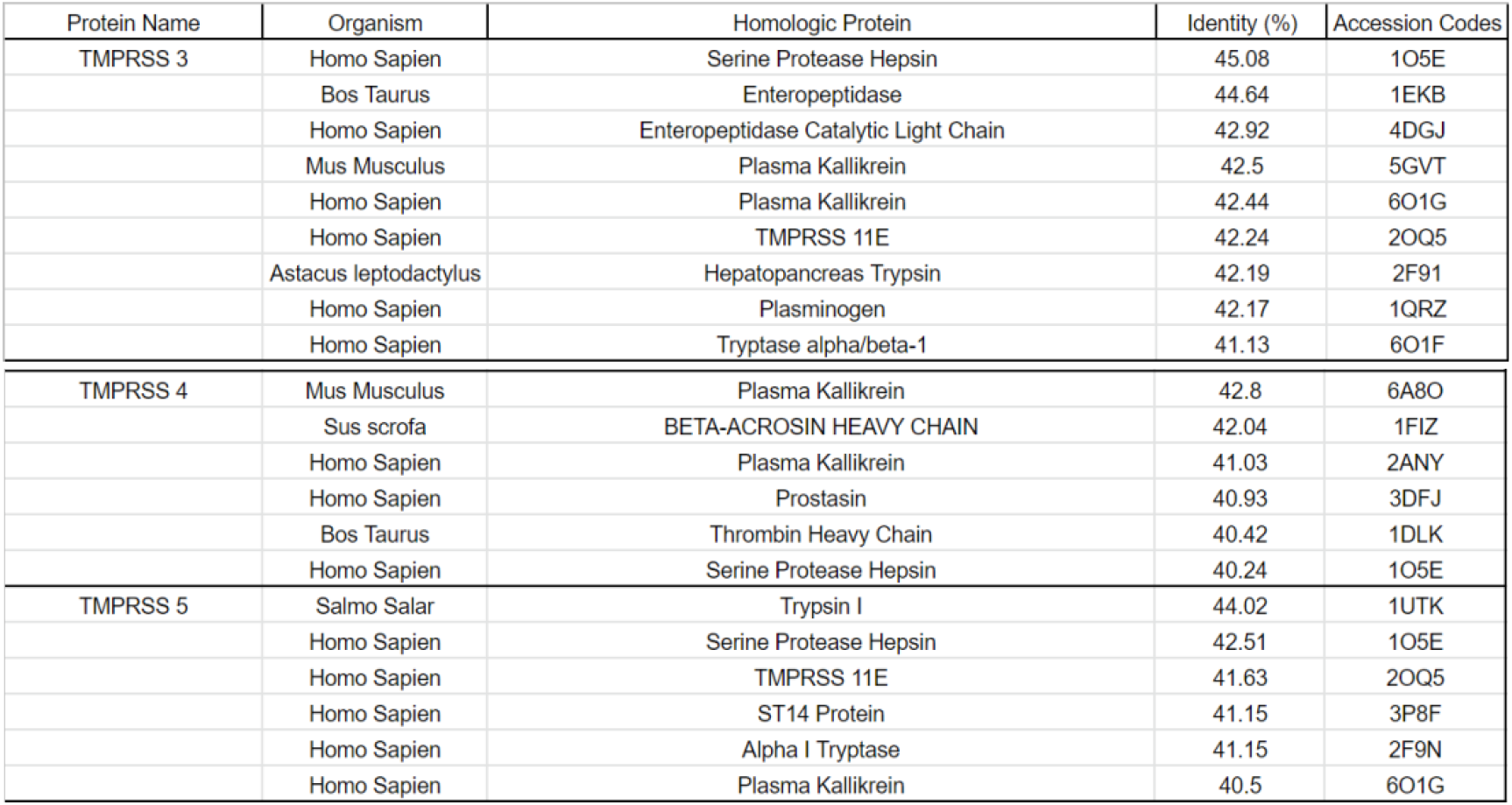
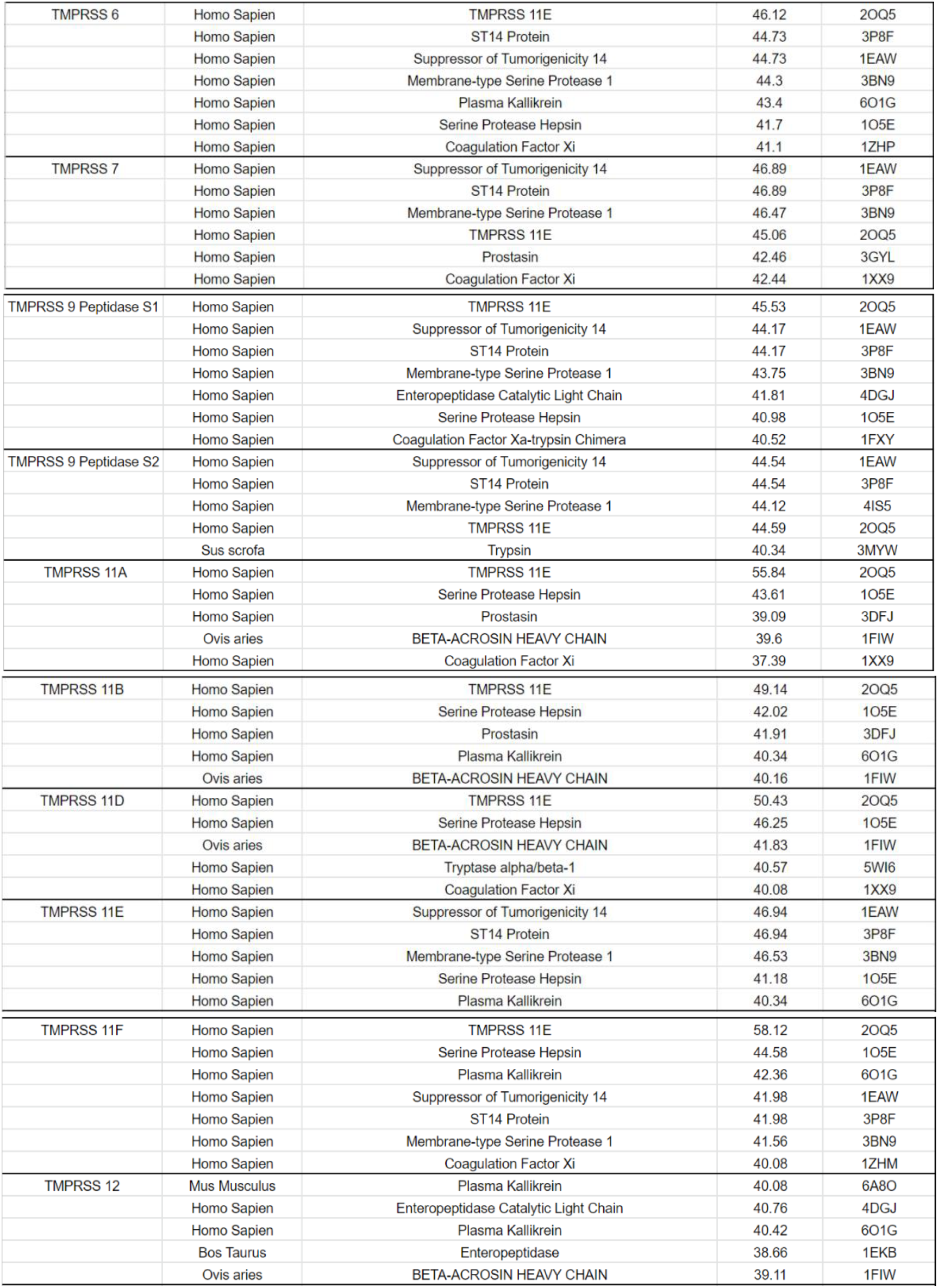
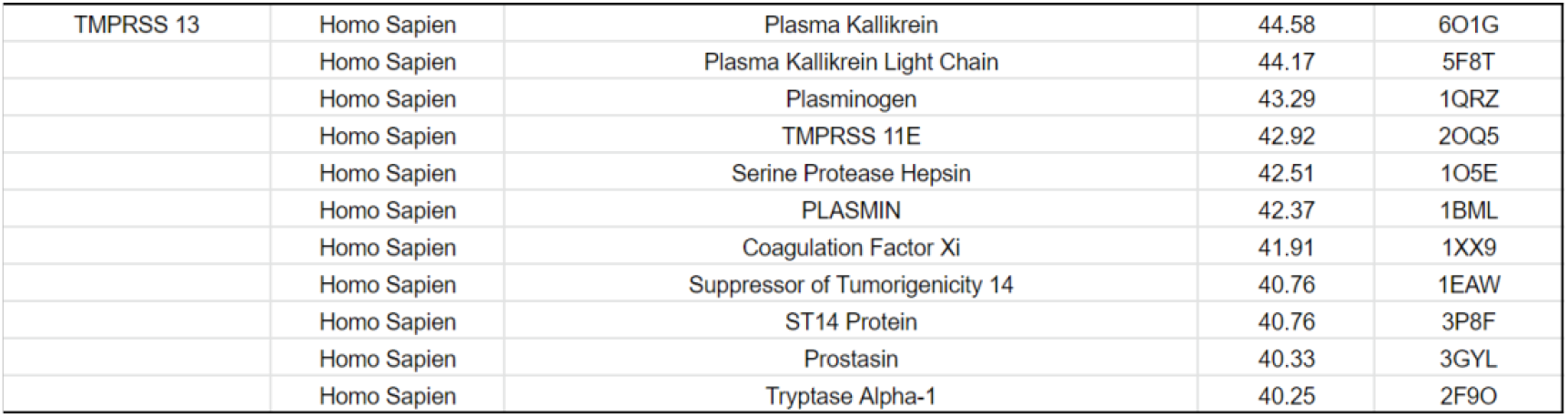
PDB codes selected with each TMPRSS enzyme.

**Supplemental Table 2.** Comparison of TMPRSS Proteins showing the catalytic triad residues, peptidase and trypsin regions. The percent identity and percent positives was calculated relative to TMPRSS2.

| Protein Name | Organism | Catalytic Triads | Peptidase Domain | Trypsin Region | Identities (%) | Positives (%) |
| --- | --- | --- | --- | --- | --- | --- |
| TMPRSS2 | Homo Sapien | H296, D345, S441 | 256-489 | 256-485 | 100 | 100 |
| TMPRSS3 | Homo Sapien | H257, D304, S401 | 217-449 | 217-445 | 53 | 65 |
| TMPRSS4 | Homo Sapien | H245, D290, S387 | 205-434 | 205-430 | 43 | 59 |
| TMPRSS5 | Homo Sapien | H258, D308, S405 | 218-453 | 218-449 | 45 | 58 |
| TMPRSS6 | Homo Sapien | H617, D668, S762 | 577-811 | 577-807 | 40 | 57 |
| TMPRSS7 | Homo Sapien | H646, D694, S790 | 606-840 | 606-836 | 40 | 56 |
| TMPRSS9 | Homo Sapien | H243, D292, S387 | 203-436 (S1) | 203-432 | 43 | 57 |
|  | Homo Sapien | H544, D592, S687 | 504-736 (S2) | 504-732 | 38 | 54 |
| TMPRSS11A | Homo Sapien | H230, D275, S371 | 190-420 | 190-416 | 39 | 54 |
| TMPRSS11B | Homo Sapien | H225, D270, S366 | 185-415 | 185-411 | 39 | 57 |
| TMPRSS11D | Homo Sapien | H227, D272, S368 | 187-417 | 187-413 | 43 | 57 |
| TMPRSS11E | Homo Sapien | H232, D277, S373 | 192-422 | 192-418 | 42 | 57 |
| TMPRSS11F | Homo Sapien | H248, D293, S389 | 206-437 | 206-433 | 42 | 60 |
| TMPRSS12 | Homo Sapien | H122, D171, S268 | 78-318 | 78-314 | 38 | 57 |
| TMPRSS13 | Homo Sapien | H366, D414, S511 | 326-559 | 326-555 | 45 | 59 |

The atomic coordinates for all the generated homology models can be found as PDB files in the supplementary .zip folder.

